# MRPL47 Deficiency Drives Mitochondrial Dysfunction via ROS/p38-MAPK/CDKN1A Signaling in Non-Small Cell Lung Cancer

**DOI:** 10.1101/2025.02.17.638626

**Authors:** Nikita Bhandari, Disha Acharya, Annesha Chatterjee, Shweta Yelshetti, Vinita Bhat, Bal Krishna Chaube, Sudhanshu Shukla

**Affiliations:** Department of Biosciences and Bioengineering, Indian Institute of Technology Dharwad, Dharwad, 580011

**Author notes:** For Correspondence Sudhanshu Shukla, Department of Biosciences and Bioengineering, Indian Institute of Technology Dharwad, Dharwad, 580011.

**Keywords:** Mitoribosome, Reactive Oxygen Species (ROS), Oxidative Phosphorylation (OXPHOS), Mitogen-Activated Protein Kinase (MAPK), CDKN1A (Cyclin-Dependent Kinase Inhibitor 1A), Cellular Senescence

## Abstract

Mitoribosomes play a pivotal role in cellular energy metabolism by synthesizing proteins involved in oxidative phosphorylation (OXPHOS) system. Dysregulation of mitoribosomes has been linked to Cancer, yet there have been no studies demonstrating the genetics and epigenetic landscape of mitoribosomal proteins (MRPs). In this study, we conducted a comprehensive analysis of expression, copy number variations, mutations, data from TCGA NSCLC patients to elucidate the genetic mechanism regulating MRPs in NSCLC. Consequently, we identified MRPL47, a significantly amplified and overexpressed mitoribosomal gene. We also found significant correlation of MRPL47 expression and copy number with patients’ survival. Functionally, we showed that inhibition of MRPL47 was associated with reduced cell proliferation and migration. Furthermore, silencing MRPL47 impaired the enzymatic activity of electron chain complex I & III leading to a defective OXPHOS system and elevated mitochondrial ROS level. Further, we showed ROS-mediated increase in CDKN1A through the p38 MAPK pathway. The increased CDKN1A level induced G1 cell cycle arrest by inhibiting E2F activity. RNA sequencing analysis further confirmed that MRPL47 hinders cell growth by inhibiting E2F pathway. Additionally, we found that MRPL47 selectively regulates mitochondrial translation of specific OXPHOS proteins rather than influencing all mitochondrial proteins. Altogether, these findings suggest that MARPL47, is amplified and overexpressed in NSCLC and plays a critical role in tumor progression by regulating ROS signaling pathways.

## 1. Introduction

Non-Small Cell Lung Cancer (NSCLC) is the most common form of lung cancer, constituting over 80% of lung cancer occurences and serving as the primary cause of cancer-related mortality worldwide ^1–3^. The discovery of novel mechanisms and biomarkers is essential for advancing NSCLC management. These findings enable early diagnosis, guide personalized therapies, and address treatment resistance. By uncovering new molecular pathways and therapeutic targets, we can develop tailored treatments for NSCLC’s heterogeneity. Novel biomarkers improve prognostic accuracy, predict treatment response, and monitor therapy efficacy in real time. Such advancements improve patient outcomes, optimize clinical decision-making, and reduce healthcare costs.

Mitochondria are essential organelles for cellular energy generation through oxidative phosphorylation. Mitoribosomes facilitate the production of proteins encoded by mitochondrial DNA that are crucial for the function of OXPHOS. ^4^. Dysregulation of mitoribosomes promotes cancer by altering mitochondrial translation and apoptosis regulation ^5^. Conversely, perturbations in oncogenic signaling pathways can impact cellular metabolism and hence influence mitochondrial activity^6^. Genetic and epigenetic modifications among cancer cells often promote a shift in metabolic pathways, switching from mitochondrial respiration to anaerobic glycolysis (Warburg effect)^7^. This transition towards the Warburg effect may be attributed to mitochondrial dysfunction along with alterations in oxidative phosphorylation activity ^8,9^. Delineating these genetic abnormalities can boost the clinical outcomes by uncovering novel therapeutic targets, early detection biomarkers, and enhanced prognostic indicators ^10–12^.

The mitoribosome, a mitochondrial-specific ribosome, synthesizes core subunits of the oxidative phosphorylation (OXPHOS) complexes essential for cellular energy production. Structurally distinct from cytoplasmic ribosomes, it is membrane-tethered and enriched with mitochondrion-specific proteins (MRPs) that enable co-translational insertion of hydrophobic OXPHOS subunits into the inner mitochondrial membrane ^13^. Dysregulation of mitoribosomal function has been increasingly linked to cancer, where altered mitochondrial metabolism contributes to tumorigenesis. For example, a pan-cancer study identified alterations in 20 MRPs associated with the early stages of mitoribosome formation ^14^. Additionally, 40 MRPs were found to be upregulated in breast cancer compared to normal tissues. Notably, MRPL52, a component of the large subunit of the mitoribosome, is a transcriptional target of HIF1α and is overexpressed in breast cancer relative to adjacent normal tissue ^15^. Mechanistically, MRPL52 helps breast cancer cells resist hypoxia-induced apoptosis. Furthermore, MRPL13 is regulated by lactate levels; its loss leads to defects in oxidative phosphorylation ^16^. MRPL33 is regulated at the splicing level by hnRNPK, and inhibition of its large variant results in reactive oxygen species (ROS) accumulation. MRPS23 serves as a crucial regulator of cell proliferation across various cancers, including breast cancer ^17^. Similarly, MRPL35 is upregulated in colorectal cancer and plays a role in regulating growth and apoptosis ^18^. These findings underscore the critical involvement of MRPs in the development and progression of multiple cancer types.

The oxidative phosphorylation (OXPHOS) complexes in mitochondria are crucial sources of reactive oxygen species (ROS), with approximately 90% of cellular ROS generated during this process. Complex I is a significant contributor to ROS production, primarily generating superoxide in the mitochondrial matrix, while Complex III is identified as the major site for ROS generation through the Q cycle^19^. Complex II also produces ROS, albeit to a lesser extent ^20^. While low levels of mitochondrial ROS play important roles in cellular signaling and metabolic adaptation, excessive ROS production can lead to oxidative damage and is implicated in various diseases, including cancer. Understanding the dynamics of ROS production at different OXPHOS complexes is essential for exploring their roles in cancer progression and potential therapeutic strategies targeting oxidative stress-related pathologies.

This study explores the role of MRPL47 in cancer cells, emphasizing its influence on mitochondrial function and tumor progression. We analyzed expression, copy number variations, and mutation data from The Cancer Genome Atlas (TCGA) in non-small cell lung cancer (NSCLC) to delineate the regulatory landscape of mitochondrial ribosomal proteins (MRPs). Our findings reveal that MRPL47 is frequently amplified and overexpressed in NSCLC. Knockdown of MRPL47 using shRNA significantly impaired cancer cell proliferation, migration, and colony formation. We demonstrated that MRPL47 selectively regulates the mitochondrial translation of specific OXPHOS proteins rather than affecting all mitochondrial proteins, leading to reduced enzymatic activity in electron transport chain complexes I and III, increased mitochondrial reactive oxygen species (ROS) levels, and a decrease in mitochondrial membrane potential (ΔΨM). Further investigations uncovered that MRPL47 modulates mitochondrial ROS signaling through the p38 mitogen-activated protein kinase (MAPK) pathway. Collectively, these findings position MRPL47 as a critical regulator of mitochondrial dynamics and ROS balance in tumor progression, suggesting new therapeutic strategies targeting this pathway.

## 2. Materials and Methods

### 2.1 Cell culture and Reagents

Human derived cell lines H1299 cells were obtained from ATCC. H460, A549 and AGS were obtained from National Centre for Cell Sciences (NCCS), Pune (India). H1299 and H460 cell lines were cultured in RPMI medium whereas, A549 and AGS cell lines were cultured in F-12 K medium supplemented with 10% fetal bovine serum (FBS), 100 U/ml Penicillin & 100 µl/ml Streptomycin (Gibco). Cell lines were maintained in a CO2 incubator at 37°C in a 5% CO_2_ humidified atmosphere.

### 2.2 Generation of stable knockdown cells

The MRPL47 gene was silenced in LUAD cell lines with the application of small interfering RNA (siRNA) or lentiviral-mediated expression of short hairpin RNA (shRNA). Western blot analysis and semi-quantitative PCR/qPCR were conducted to assess the efficiency of MRPL47 silencing. For shRNA-mediated knockdown, two distinct shRNAs were employed for MRPL47 silencing: sh1 CTTGCCTTATGTGGACCATTT and sh2 GTAGATTCCATGGATGCATTA .pLKO.1 puro, a 3rd generation lentiviral plasmid, for the cloning and silencing of MRPL47 expression in LUAD cell lines was acquired from Addgene (Plasmid #8453). This was accomplished by designing a 21 bp target sequence for MRPL47 and subsequently cloning it into the Age1-EcoR1 sites of the pLKO vector. The Lenti-X 293T cell line was utilized for the generation of viral supernatants and subsequent infections. Transfection was performed using Xfect polymer (TaKaRa Bio, cat.#631317) in accordance with the manufacturer’s guidelines. The transfected cells were selected with 2.5 µg/mL puromycin (Sigma, cat.#P8833) for at least 48 hours. For siRNA mediated RNA interference, cells were transfected with 5 nM Ambion™ Silencer™ Negative Control #1 siRNA (siNT) (Thermo Fisher Scientific, cat #AM4611) and MRPL47 siRNA (siMRPL47) utilizing Lipofectamine™ RNAiMAX transfection reagent (Thermo Fisher Scientific, cat #13778100), in accordance with the manufacturer’s guidelines. The employed target sequence was as follows: siMRPL47 GACAUCUUUGGAAGAAUCA. Finally, these stable knockdown cells were harvested for protein and RNA extraction.

### 2.3 Semi-quantitative/Quantitative real-time PCR

Total RNA was extracted utilizing TRIzol reagent (Ambion, cat.#15596018), and 500 ng of total RNA was employed to synthesize cDNA with iScript RT mix (Biorad, cat.#1708840), according to the manufacturer’s instructions followed by a semi-quantitative PCR or a quantitative real time PCR employing Emerald GT PCR mix (TaKaRa, cat.#RR310A) and universal SYBR Green mix (Biorad, cat#1725271) in a thermal cycler.The ΔCt method was employed to quantify the mRNA levels of the target gene,normalised against TPT1, housekeeping gene. The 2^-ΔΔCt^ technique was employed to compare the mRNA levels of each target gene, and the relative amplification values were illustrated in the graph. The following primer sequences were used for MRPL47_FW: 5′-TGGAAGAATCATCTGGCACA-3′; MRPL47_RV: 5′-ACTTTTGGGCTTCAGCAAGA-3′ TPT1_FW: 5′-GATCGCGGACGGGTTGT-3′ TPT1_RV: 5′-TTCAGCGGAGGCATTTCC-3′

### 2.4 Cell proliferation assay

To assess cell viability and proliferation rate post-knockdown, we conducted an MTT tassay utilizing yellow tetrazolium MTT [3-(4,5-dimethylthiazol-2-yl)-2,5-diphenyl-tetrazolium bromide] (Sigma, cat. #475989). 20,000 cells per well were seeded in a 96-well plate and incubated with MTT for 4 hours in a CO2 incubator followed by the addition of DMSO to solubilize the purple formazan complex, thereafter measuring the absorbance at 570 nm with a spectrophotometer. For each cell type, the linear correlation between cell count and signal output was determined, enabling the quantification of variations in the rate of cell proliferation.

### 2.5 Wound healing Assay

The migration of cells was assessed using a scratch wound assay performed on the Incucyte® Live-Cell Analysis System (Sartorius) equipped with the Scratch Wound module. Cells were seeded into an Incucyte imagelock 96-well plate (Sartorius, cat. #BA-04856) at a density of 50,000 cells/well. Subsequently, uniform scratches were created in the cell monolayer using the Incucyte® WoundMaker™ tool. Images of the wound area were automatically captured at an interval of 4 hours over a period of 24 hours using the Incucyte® system. The wound width and confluence were quantified using the Incucyte® Scratch Wound module, which calculates cell migration based on the decrease in the wound area over time. Data were analyzed using Incucyte® software and are presented as a percentage of wound closure relative to baseline. Finally, the wound healing % vs Hours graph was plotted using Graphpad Prism software.

### 2.6 Colony Suppression Assay

To identify the ability of knockdown cells to grow into colonies, we seeded 500 cells/well in a 12-well plate and allowed them to grow for 2 weeks. The media was replaced every 4-5 days for 12 days followed by staining the colonies using 0.25% crystal violet solution. Finally, the images were captured and analyzed.

### 2.7 Western Blot Analysis

To identify the changes in protein expression of the knockdown cells, western blot was performed using MRPL47 antibody. For protein isolation, cells were trypsinized and washed with cold PBS followed by addition of IP lysis buffer (Pierce #87787) and protease inhibitor cocktail (Halt protease inhibitor cocktail #87786) to the pellet. The cell lysates were centrifuged at 13000 g for 15 minutes and supernatants were collected. The total protein concentration was estimated before loading onto polyacrylamide gel by BCA protein assay kit (Pierce #23227) and a total of 50 µg of cell lysate was loaded and resolved in 12% polyacrylamide gel (Biorad TGX FastCast Acrylamide Kit 12% 161-0175)). The gel was blotted in PVDF membrane (Immun-Blot PVDF Membranes for Protein Blotting, Biorad #1620177) and the membrane was blocked using a 5% blocker solution (Blotting Grade Blocker, Biorad # 1706404). The membrane was incubated with MRPL47 (Immunotag #ITT03673) and internal control Actin B (ABclonal #AC004) or GAPDH (ABclonal #AC002) antibodies were used. The membrane was later incubated with HRP-tagged goat anti-mouse (Bio Rad #172-1011) or anti-rabbit (Invitrogen #31460) secondary antibodies. All the steps were followed by washing with TBS (Bio Rad #1706435) comprising 0.1% Tween 20 (Polysorbate 20, MP Biomedicals #103168). The bound antibody complexes were visualized using ECL Western Blotting substrate (Pierce #32209) or femtoLUCENT PLUS HRP chemiluminescent reagents (G Biosciences #786-003).

### 2.8 Determination of Mitochondrial ROS production

The intracellular mitochondrial ROS level was determined using MitoSOX red mitochondrial superoxide indicator (Invitrogen #M36008). 0.3 x 10^6^ cells were seeded in a 6 well plate and kept for growing inside the CO2 incubator overnight. Next day cells were treated with 2 mM MitoSOX, mitochondrial superoxide indicator for 10 min. in a CO2 incubator at 37°C & 5% CO2. The treated samples were washed three times with PBS. The cells were then trypsinized and collected in FACS tubes. Unstained & unstimulated cells were used as a control while performing FACS in atteuneNXT Flow cytometer (Thermofisher). Finally, the fluorescence was recorded at 510/580 nm and the results were analyzed in FCS express software.

### 2.9 Senescence Assay

To identify the senescence in MRPL47 knockdown cells, Beta-galactosidase staining kit (CST #9860) was used. Control and experimental cells were allowed to continuously grow in a 6 well plate for the experiment for one week prior to the experiment to achieve senescence. Cells were then fixed with 1x fixating solution for 10-15 min at RT. Next, PBS wash was performed two times followed by addition of Beta galactosidase staining solution. The plate was then sealed tightly with a parafilm and incubated in a dry CO2 free incubator overnight. Next day the image was acquired using an optical microscope.

### 2.10 Extracellular Oxygen Consumption Assay

Changes in Oxygen consumption after knocking down MRPL47 in H460 cells was measured using commercial kit (abcam #ab197243). In this assay an oil layer was added on top of the assay medium to limit the oxygen diffusion. The fluorescent dye was quenched by oxygen present in the medium. Finally real time kinetics analysis was performed using fluorescent based reader.

### 2.11 SuNSET Assay

To detect the changes in the newly synthesized protein synthesis, SUrface SEnsing of Translation (SUnSET) assay was performed that measures the puromycin incorporated into nascent polypeptide peptide chains using anti-puromycin antibody. After silencing MRPL47, the cells were treated with 1 μM Puromycin (Sigma #P8833) for 30 min and incubated at 37°C. Post treatment, the mitochondria were isolated using manufacturer’s protocol using a mitochondrial isolation kit (Abclonal #Ab110170). Furthermore, 50 μg of mitochondrial extract was resolved using a 12% polyacrylamide gel. Following this the samples were transferred to a PVDF membrane (G Biosciences #GS-PVDF-302) and subsequently incubated in TBST buffer consisting of 5% Quick Blocker protein powder (G Biosciences #786–011) for two hours at room temperature. Next, PVDF membrane was cut and incubated overnight at 4°C with anti-Puromycin (DSHB #PMY-2A4), MRPL47 (Immunotag #ITT03673), GAPDH (ABclonal #AC002) antibodies. Following day, membranes were incubated for 1 Hour at room temperature in TBST buffer containing HRP tagged goat anti-mouse (Bio Rad #172–1011) or anti-rabbit (Invitrogen #31,460) secondary antibodies. All the membrane washing steps were performed four times at an interval of 10 min using TBS (HIMEDIA #ML029) along with 0.1% Polysorbate 20 (MP Biomedicals #103,168). Finally, to develop the image of the blot containing protein of interest, chemiluminescence detection method was performed using an ECL Western Blotting substrate (Pierce #32,209) or femto LUCENT PLUSHRP (G Biosciences #786–003) reagents.

## 3. Results

### 3.1. Genomic alterations in MRPL47 reveal Its oncogenic potential in NSCLC

Genomic alterations in MRPL47 reveal its oncogenic potential in non-small cell lung cancer (NSCLC). The structural organization of the 55S mammalian mitoribosome consists of two distinct subunits: a large 39S subunit and a small 28S subunit ^21,22^. This complex molecular machine is uniquely adapted for mitochondrial translation, comprising 82 nuclear-encoded mitochondrial ribosomal proteins (MRPs) alongside mitochondrial-encoded RNA components ^23^ (**Figure 1A**). To investigate the genetic aberrations associated with MRP genes in NSCLC, we analyzed genomic and transcriptomic data from The Cancer Genome Atlas (TCGA). Our analysis revealed that five MRP genes—MRPL47, MRPL36, MRPL9, MRPS30, and MRPS10—exhibited a high frequency of genomic alterations (>10%), with MRPL47 displaying the highest frequency (**Figure 1B**). Notably, we did not identify any recurrent mutations in these MRP genes, indicating that the wild-type function of MRPs is crucial for NSCLC growth. Using cBioPortal, we found that all five genes are significantly amplified in NSCLC patients, with a strong correlation to their expression levels (**Figure 1C**). MRPL47, being the most amplified MRP gene, showed a robust linear correlation between expression and copy number variation (CNV) (**Figure 1D**). Consistent with this, an independent TCGA dataset validated that copy number amplification drives MRPL47 overexpression in lung cancer, with amplified samples exhibiting significantly elevated MRPL47 expression compared to normal and diploid samples (p < 0.0001) (**Figure 1E**). The immunohistochemistry data from human protein atlas also showed increased expression of MRPL47 protein in human lung cancer compared to normal (**Figure 1F and G**). Kaplan-Meier survival analysis indicated that high MRPL47 expression is associated with significantly poorer overall survival in both microarray and TCGA LUAD datasets (p < 0.0001), underscoring its prognostic significance in LUAD patients (**Figure 1H**). We also used UCSC genome browser to show thar MRPL47 highly conserved gene with high promoter activity (**Supplementary Figure 1A**). Collectively, these observations suggest that MRPL47 is overexpressed due to recurrent genetic amplification in NSCLC. We analyzed the expression and amplification levels of MRPL47 across various cancer types and found it to be consistently amplified and overexpressed in all examined cancers (**Supplementary Figure 1B, C, and D).** Moreover, MRPL47 expression demonstrated high specificity and sensitivity in distinguishing cancerous from normal tissues. Additionally, elevated MRPL47 expression was associated with poor survival outcomes in colorectal, pancreatic, breast, and ovarian cancers (**Supplementary Figure 1E, F, and G**).

**Figure 1.**
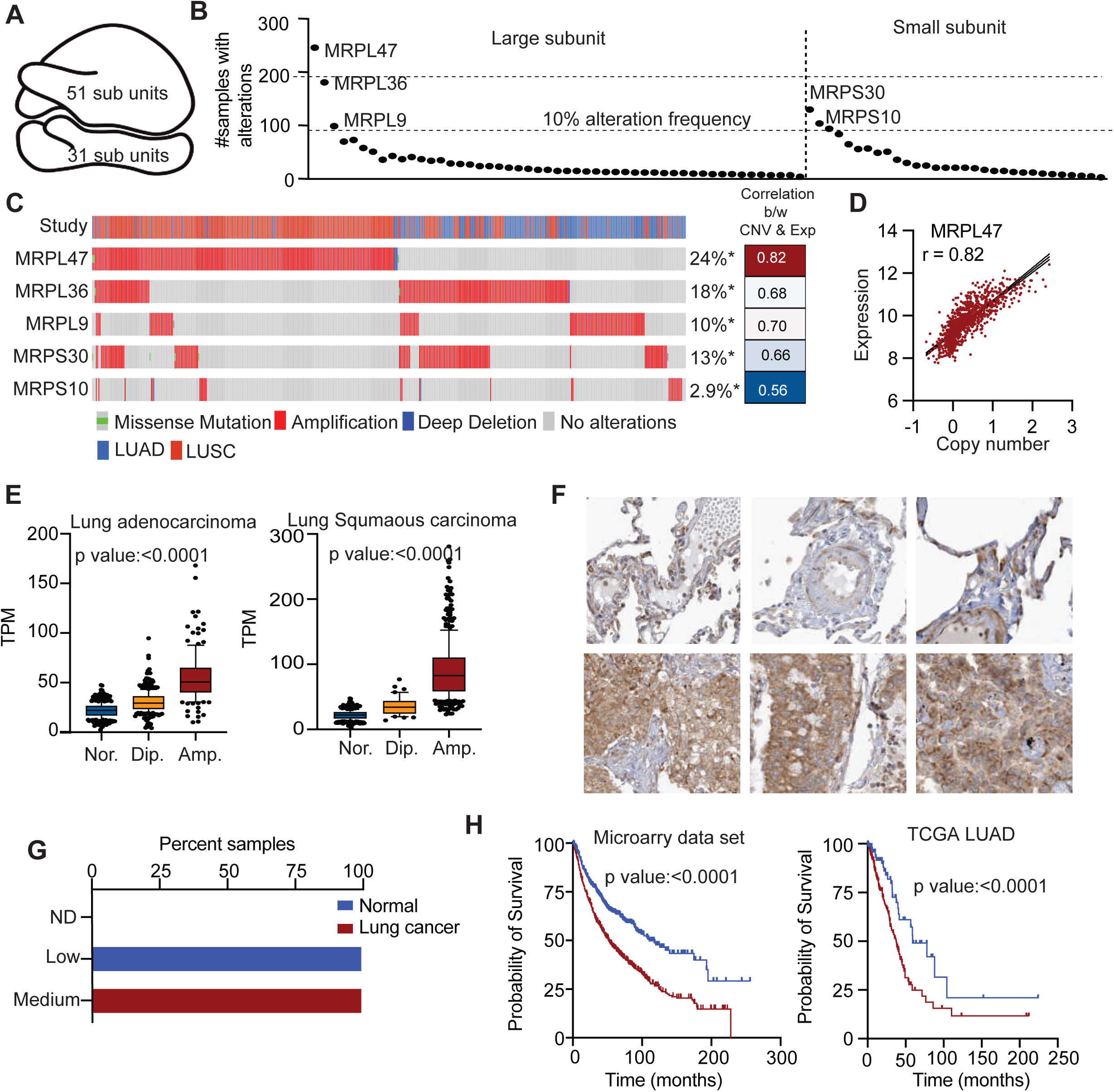
Amplification and Overexpression of MRPL47 contributes to reduced overall survival in non-small cell lung cancer (NSCLC). **(A)** Structural organization of a human mitoribosome consisting of a lager and a smaller subunit, essential for mitochondrial translation. **(B)** The genomic analysis of top five MRP genes across NSCLC samples exhibiting an alteration frequency >10%, with MRPL47 displaying the highest alteration rate. **(C)** The cBioPortal analysis highlights distinct genetic alterations in candidate MRP genes among NSCLC patients, with the frequency of gene alterations represented as percentages. MRPL47 exhibited the highest alteration frequency (24%), primarily driven by amplifications. **(D)** Graph depicting a positive correlation between copy number variations (CNVs) and gene expression for the five identified MRP genes, with MRPL47 exhibiting the strongest correlation (r = 0.82). **(E)** MRPL47 expression levels categorized by CNV status in LUAD and LUSC subtypes. Amplified (Amp.) samples displayed .significantly higher expression levels than normal (Nor.) or diploid (Dip.) samples (p < 0.0001). **(F)** Immunohistochemistry (IHC) analysis of MRPL47 expression in lung cancer and normal tissue. Lung cancer samples exhibit stronger MRPL47 staining (bottom panel) compared to normal lung tissue (top panel), indicating increased MRPL47 protein expression in tumor cells.**(G)** Bar graph indicating the distribution of MRPL47 expression levels across normal and lung cancer samples. Lung cancer samples predominantly exhibit medium to high MRPL47 expression, whereas normal samples show lower or undetectable expression, suggesting MRPL47 upregulation in lung cancer **(H)** Kaplan-Meier survival analysis showing a correlation between high MRPL47 expression and poor overall survival in both microarray and TCGA LUAD datasets (p < 0.0001).

### 3.2 MRPL47 is required for the cell proliferation and migration

To investigate the impact of MRPL47 on cellular growth, two LUAD cell lines, A549 and H1299, were selected for the experiments. Two distinct shRNAs targeting MRPL47 RNA were used for knockdown, which was subsequently validated by analyzing the protein expression of A549 cells **(Figure 2A)**. Interestingly, we observed a decrease in the proliferation of A549 MRPL47 knockdown **(Figure 2B, G)**. Colony suppression assay also showed that MRPL47 knockdown cells show reduced long-term cell growth in A549 cell line **(Figure 2C)**. Next, we performed a wound-healing assay displaying a reduced wound healing as compared to the control cells **(Figure 2D).** These observations were replicated in H1299 cells **(Figure F-G).** In conclusion, MRPL47 is essential for cancer cell growth and proliferation through its role in mitochondrial function. Silencing of MRPL47 in LUAD cells inhibited proliferation, growth, and migration, highlighting its potential as a therapeutic target.

**Figure 2.**
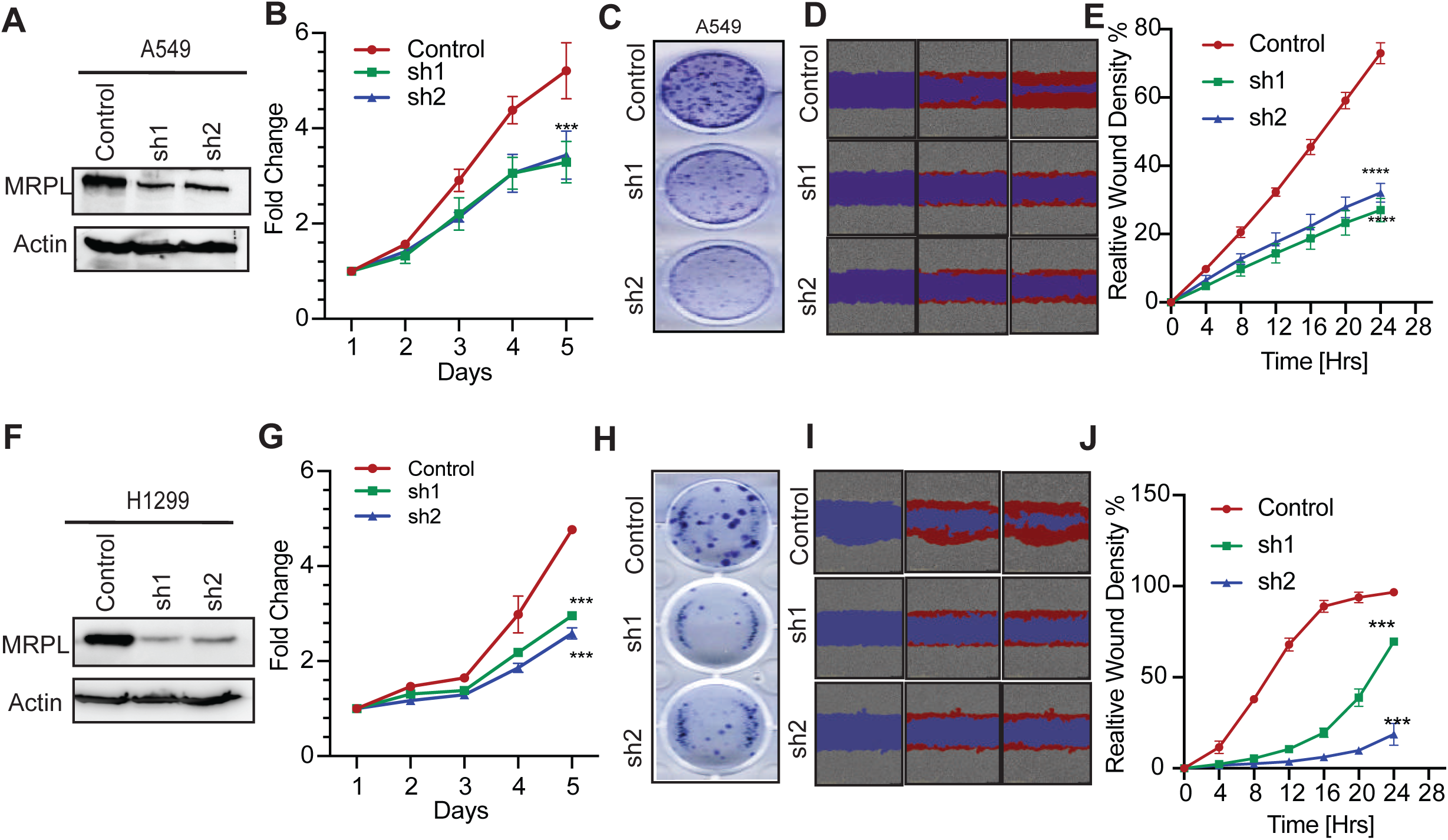
MRPL47 knockdown suppresses cell proliferation, colony formation, and migration in LUAD cell lines A549 and H1299. **(A, F)** Western blot analysis confirming MRPL47 knockdown using two independent shRNAs) in A549 (A) and H1299 (F) cells. **(B, G)** Cell proliferation assay demonstrating reduced proliferation in MRPL47 silenced cells compared to control in A549 (B) and H1299 (G) cells over a 5-day period. **(C, H)** Colony formation assay showing decreased colony-forming ability in MRPL47 knockdown cells in A549 (C) and H1299 (H) cell lines. **(D, I)** Representative images of Scratch wound-healing assay displaying delayed wound closure in MRPL47 silenced cells compared to controls in A549 (D) and H1299 (I) cells **(E, J)** Quantification of wound-healing assay indicating significantly reduced migration capacity in MRPL47 depleted cells in A549 (E) and H1299 (J) cells over a time period of 24 hours. *Statistical significance:* (***denotes p < 0.001; ****denotes p < 0.0001). Error bars represent mean ± SD from triplicate experiments.

### 3.3 MRPL47 is not a universal mitoribosome subunit; rather, it specifically regulates the translation of certain mitochondrially encoded proteins

Previous studies have identified that knocking down MRPs such as MRPL12 and MRPS34, leads to a global reduction in the mitochondrial translation, highlighting their critical role in regulating mitochondrial protein synthesis ^24,25^. Based on these findings, we aimed to investigate the role of MRPL47 in mitochondrial translation beginning with an assessment of its localization. The western blot results demonstrated the predominant localization of MRPL47 in the mitochondrial fraction of H460 cell line **(Figure 3A)**. Next, we examined the levels of COX1, a mitochondrially encoded protein in MRPL47 silenced cells. Surprisingly, MRPL47 knockdown did not affect COX1 expression, whereas a clear reduction was observed in the linezolid-treated samples confirming impaired mitochondrial translation **(Figure 3B)**. Motivated by these results, we sought to determine whether MRPL47 is required for the synthesis of global mitochondrial translation or its regulation is limited to translation of specific mitochondrially-encoded proteins. To address this question, we performed a SUnSET (Surface SEnsing of Translation) assay for measuring global protein synthesis in MRPL47 silenced cells. This assay utilizes puromycin’s ability to incorporate into the newly synthesized polypeptide chains, prematurely terminating translation. These puromycin labelled proteins can be further detected by using an anti-puromycin antibody. Our findings revealed a robust puromycin incorporation signal across multiple molecular weight bands, indicating an active global protein synthesis in the control cells. In contrast, MRPL47 knockdown cells exhibited a significant reduction in the puromycin signal of some proteins, but not all, validating our previous findings. **(Figure 3C)**. To further explore the involvement of MRPL47 in the functionality of OXPHOS system we utilized a total OXPHOS monoclonal antibody cocktail targeting multiple subunits of the respiratory chain complex crucial for oxidative phosphorylation. We observed a reduction in the Complex I (NDUFB8), II (SDHB), and IV (COXII) subunits while Complex III (UQCRC2) and Complex V (ATP5A) expression remained unchanged indicating selective impairment in the respiratory complexes, consistent with our previous findings. **(Figure 3D)**. This selective disruption within the electron transport chain subunits suggests specific dysfunctions such as a deficiency of NDUFB8 subunit is known to diminish electron flow and ATP production whereas, SDHB deficiency leads to abnormal succinate accumulation further inducing tumorigenesis ^26^. Similarly, COXII subunit is involved in regulation of the final step of ETC, where oxygen is utilized to produce water ^27^. These findings suggest that MRPL47 silencing selectively impacts mitochondrial protein synthesis and the functionality of the OXPHOS system, thereby affecting cellular energy metabolism.

**Figure 3.**
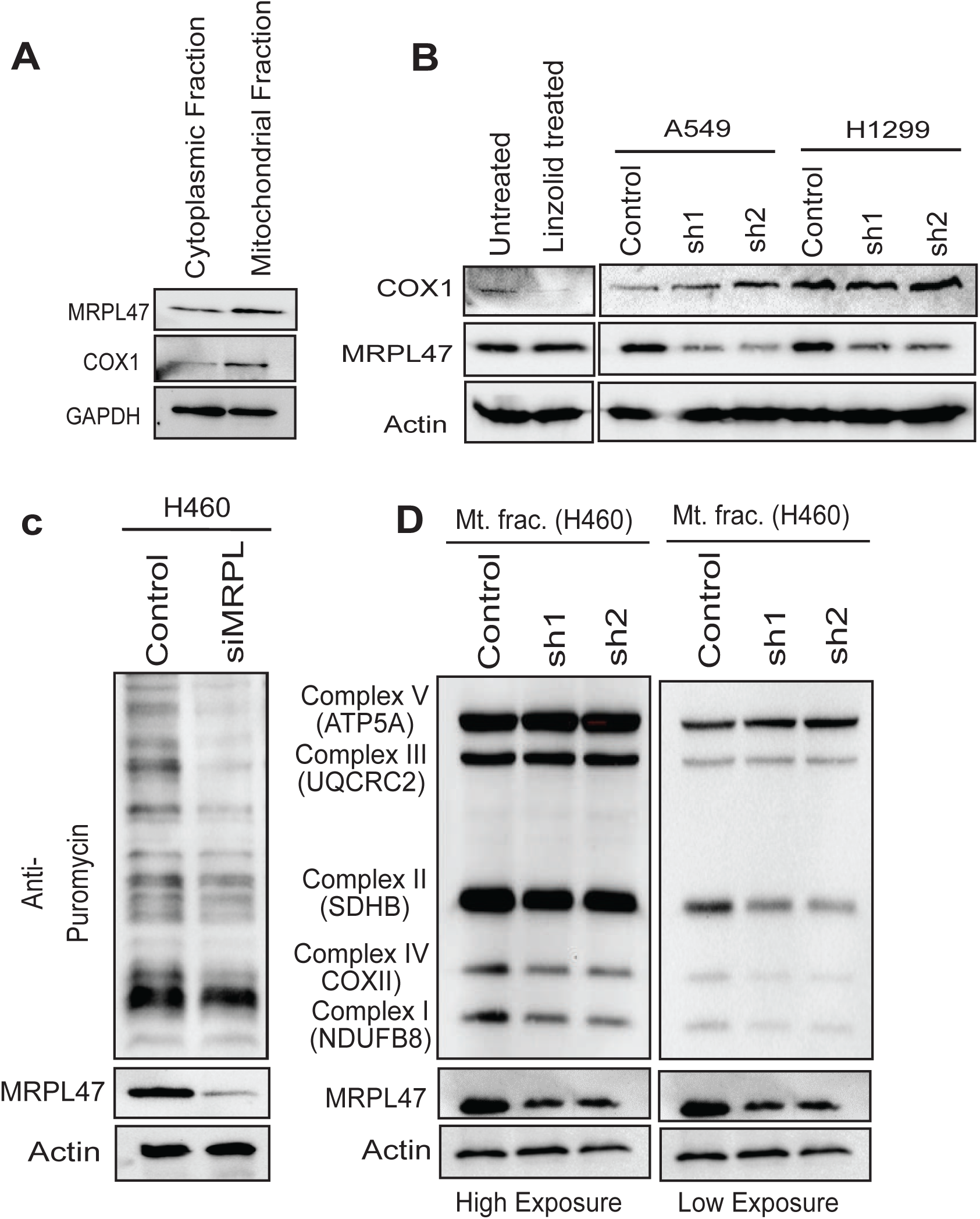
Role of MRPL47 in Mitochondrial Function and Respiratory Complex Activity. **(A)** Western blot analysis showing MRPL47 localization predominantly in the mitochondrial fraction, alongside COX1 (mitochondrial marker) and GAPDH (cytoplasmic marker) in H460 cells. **(B)** COX1 protein levels in A549 and H1299 cells showing unchanged expression in control vs knockdown cells. A decrease in the COX1 level of linzolid treated cells indicate inhibition of mitochondrial translation. **(C)** SUnSET assay demonstrating a significant decrease in some but not all mitochondrial protein translation upon MRPL47 knockdown in H460 cells, confirming MRPL47’s role in selective mitochondrial translation. **(E)** Western blot analysis of mitochondrial respiratory complex subunits (Complexes I-V) in mitochondrial fractions from H460 cells. MRPL47 knockdown results in reduced levels of Complex I (NDUFB8), Complex II (SDHB), and Complex IV (COXII), while Complex III (UQCRC2) and Complex V (ATP5A) remain unaffected.

### 3.4 Mitoribosomal dysfunction impairs OXPHOS further leading to ROS generation

The biogenesis of Mitoribosome is crucial for regulation of mitochondrial respiration and cellular energy homeostasis. Often, mitoribosome dysfunction can contribute to impairment of the OXPHOS system which in turn leads to elevated ROS production causing detrimental effects on cellular function. ^28^. To elucidate the function of MRPL47 in mitochondrial respiration, we measured the OCR (Oxygen Consumption Rate) in MRPL47 silenced cells using high-resolution respirometry with the Oroboros O2K system. The oxygraph indicated a progressive decrease in the Baseline oxygen consumption over time in the MRPL47 silenced cells when compared to the control, while compound-specific injections (ce1-ce4) demonstrated differential effects on mitochondrial activity such as CE1 induced respiration, indicating diminished basal respiration in MRPL47-silenced cells. CE2 (Oligomycin) inhibited Complex I, hence validating the disruption of the respiratory chain. CE3 (FCCP), an uncoupler, elicited maximal electron flux, demonstrating an impaired oxidative phosphorylation due to the reduced response. CE4 (Antimycin A) completely obstructed respiration, emphasizing the dependency of mitochondria on OCR and the bioenergetic deficiencies resulting from MRPL47 ablation. **(Figure 4A)**. Furthermore, OCR profiles displayed a significant reduction across basal, ATP-linked, maximal as well as non-mitochondrial respiration states strongly suggesting the critical role of MRPL4 in regulating mitochondrial respiration and OXPHOS activity. **(Figure 4B).** Upon sequential treatment with the inhibitors-oligomycin, FCCP, and antimycin A, control cells displayed a higher baseline and maximal respiration, while knockdown cells maintained consistently lower OCR, reinforcing mitochondrial dysfunction as a specific consequence of MRPL47 loss **(Figure 4C).** The observed changes in OCR profiles, particularly reductions in ATP-linked and maximal respiration, suggest impaired ETC activity, leading to electron leakage and ROS generation. To validate this, flow cytometry analysis was performed using MitoSOX, a mitochondrial superoxide indicator showing elevated ROS levels in knockdown cells compared to the control **(Figure 4D and E).** Since, ETC complex I and III are considered major sites for ROS production, we aimed to perform a spectrophotometric assessment of enzymatic activities of these complexes. Complex I dysfunction, evidenced by reduced NADH oxidation, aligns with decreased NDUFB8 protein levels in MRPL47 knockdown cells, explaining the diminished OCR observed in the Oroboros O2k assay. Complex III dysfunction, despite unchanged UQCRC2 protein levels, reflects impaired enzymatic activity likely due to ETC destabilization from upstream defects in Complex I **(Figure 4F,G)**. Furthermore, elevated ROS level disrupts the ETC, leading to a reduction in the proton gradient and mitochondrial membrane potential. Consistently, TMRM staining revealed a marked reduction in the fluorescence intensity of the knockdown cells, confirming impaired mitochondrial membrane potential and dysfunction upon MRPL47 silencing **(Figure 4H).** According to our previous findings, silencing MRPL47 significantly reduced cell proliferation. To determine whether this effect was due to oxidative stress or ROS, we performed a cell proliferation assay by treating the cells with N-acetyl-L-cysteine (NAC), a well-known ROS scavenger. Treatment with NAC rescued the cell proliferation deficit, indicating the observed reduction in cell proliferation as a result elevated ROS levels **(Figure 4I).** Since, impaired membrane potential also affects the proton gradient essential for ATP synthase activity, we further validated this through a luminescence assay showing significantly decreased ATP generation over a time period of 5 hrs. in the knockdown groups. **(Figure 4J).** Collectively, these results demonstrate that MRPL47 plays a pivotal role in mitochondrial function by regulating respiration, oxidative stress, membrane potential, and energy production. Its silencing leads to mitochondrial dysfunction and impaired cellular proliferation, largely driven by increased ROS levels.

**Figure 4.**
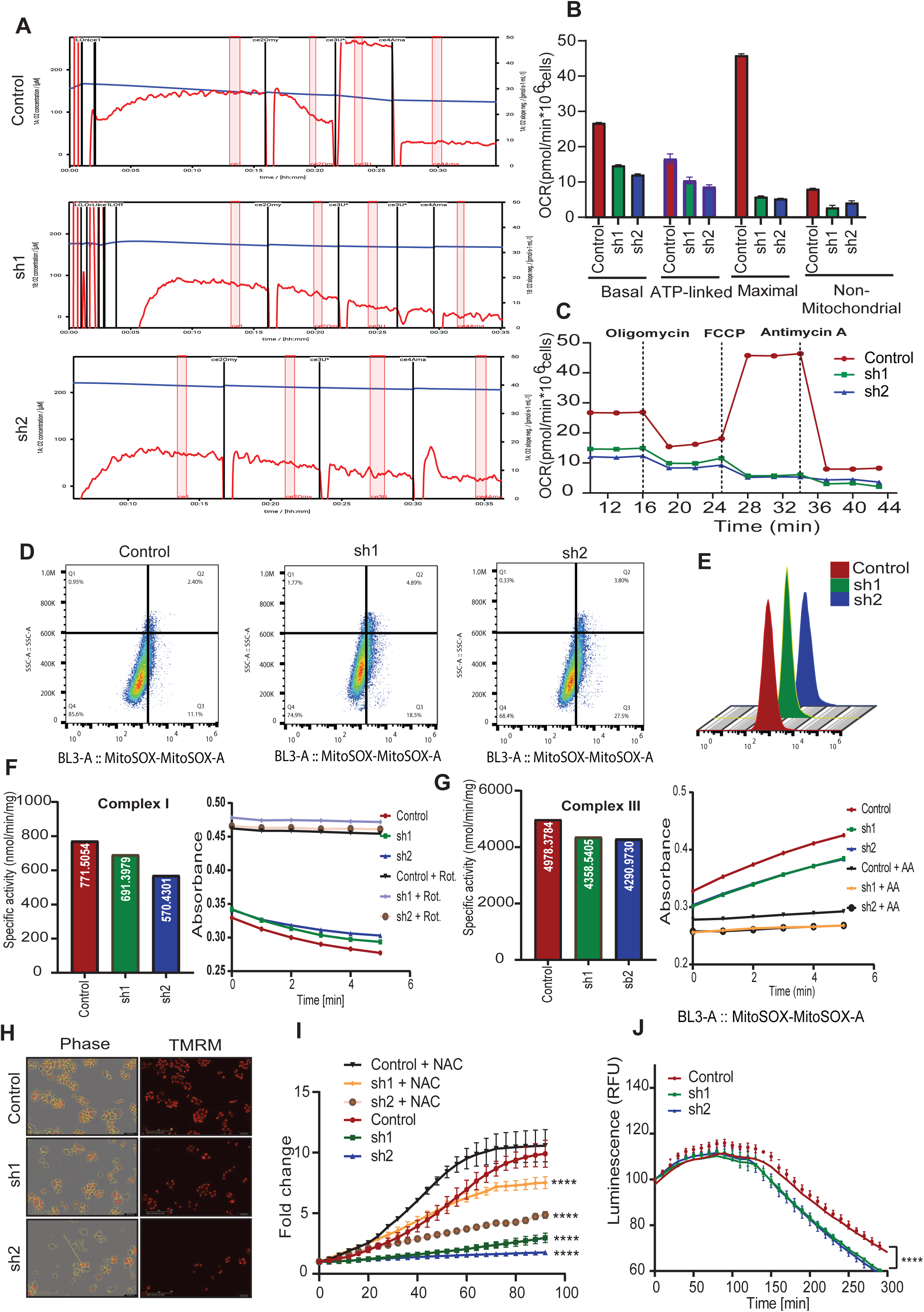
Functional consequences of MRPL47 gene silencing on mitochondrial dynamics, oxidative stress, and cell growth. **(A)** Assessment of mitochondrial bioenergetics in H460 cells using high-resolution respirometry. Oroboros O2k representative oxygen consumption rate (OCR) tracings of control versus MRPL47 knockdown groups. Measurements recorded under basal conditions, following sequential injections of inhibitors-oligomycin (ATP synthase inhibitor), FCCP (uncoupler), and antimycin A (complex III inhibitor). **(B)** Quantification of basal respiration, ATP-linked respiration, maximal respiration, and non-mitochondrial respiration based on OCR measurements. **(C)** Time-course representation of OCR changes in response to mitochondrial inhibitors in control and MRPL47 knockdown cells, demonstrating a substantial decrease in mitochondrial respiratory capacity in knockdown groups. **(D)** MitoSOX-based flow cytometric detection of mitochondrial ROS production in control and MRPL47 deficient H460 cells Increased MitoSOX fluorescence corresponds to the higher levels of mitochondrial oxidative stress in MRPL47 knockdown cells. **(E)** Representative flow cytometry histogram overlays displaying MitoSox Red fluorescence intensity in control, sh1, and sh2 cells. MRPL47 knockdown cells exhibit a rightward shift, confirming elevated mitochondrial oxidative stress. **(F,G)** Enzymatic activity of mitochondrial Complex I and III measured via spectrophotometry. Specific activity (left) and absorbance over time (right) show a reduction in activity for both complexes in MRPL47 knockdown cells. Addition of specific inhibitors (Rotenone for Complex I, Antimycin A for Complex III) confirms specificity of the assays. **(H)** Incucyte images showing phase contrast and Tetramethylrhodamine methyl ester (TMRM) staining images of H460 cells to evaluate changes in mitochondrial membrane potential (ΔΨm). Loss of TMRM fluorescence signal in MRPL47 knockdown cells suggests depolarization of the mitochondrial membrane. **(I)** Rescue experiment validating ROS mediated inhibition of cell proliferation in MRPL47 silenced cells, with or without NAC treatment. Elevated ROS levels in sh1 and sh2 cells are rescued by NAC treatment. **(J)** Luminescence-based ATP assay showing a significant decline in ATP generation over a time duration of 5 hrs in control and MRPL47 knockdown cells. A marked reduction in ATP production is observed in sh1 and sh2 groups, reflecting impaired mitochondrial function.

### 3.5. Silencing of MRPL47 inhibits E2F pathway

To understand the molecular consequences of MRPL47 mediated cell growth inhibition, we performed RNA-seq analysis to identify differentially expressed genes (DEGs) in MRPL47-silenced cells compared to controls. The heat map representation of DEGs indicated substantial changes in gene expression of MRPL47-silenced genes relative to the controls **(Figure 5A)**. Gene ontology (GO) enrichment analysis using down regulated genes highlighted enrichment of pathways related to cell cycle progression, chromosome maintenance and DNA unwinding, highlighting a functional link between MRPL47 and genomic integrity **(Figure 5B)**. Further analysis of specific gene sets confirmed the downregulation of key genes involved in cell cycle, chromosome maintenance and DNA unwinding, suggesting defects in DNA replication and genomic stability **(Figure 5C)**. Further detailed analysis showed the downregulation of various positive E2F targets in MRPL47 silenced cells. Whereas many genes negative regulated by E2F, were downregulated in MRPL47 knockdown cells **(Figure 5D)**. Pathway enrichment analysis validated these findings, through illustrating significant enrichment of E2F target genes in of cell cycle checkpoint and replication stress response pathways, suggesting the role of MRPL47 in replication stress and growth arrest **(Figure 5E).** Further, we showed that many E2F target genes associated with cell cycle regulation and replication were upregulated **(Figure 5F).** Altogether, these findings demonstrate that loss of MRPL47 severely impacts cell cycle regulation, DNA replication chromosomal integrity by inhibiting E2F mediated transcription.

**Figure 5.**
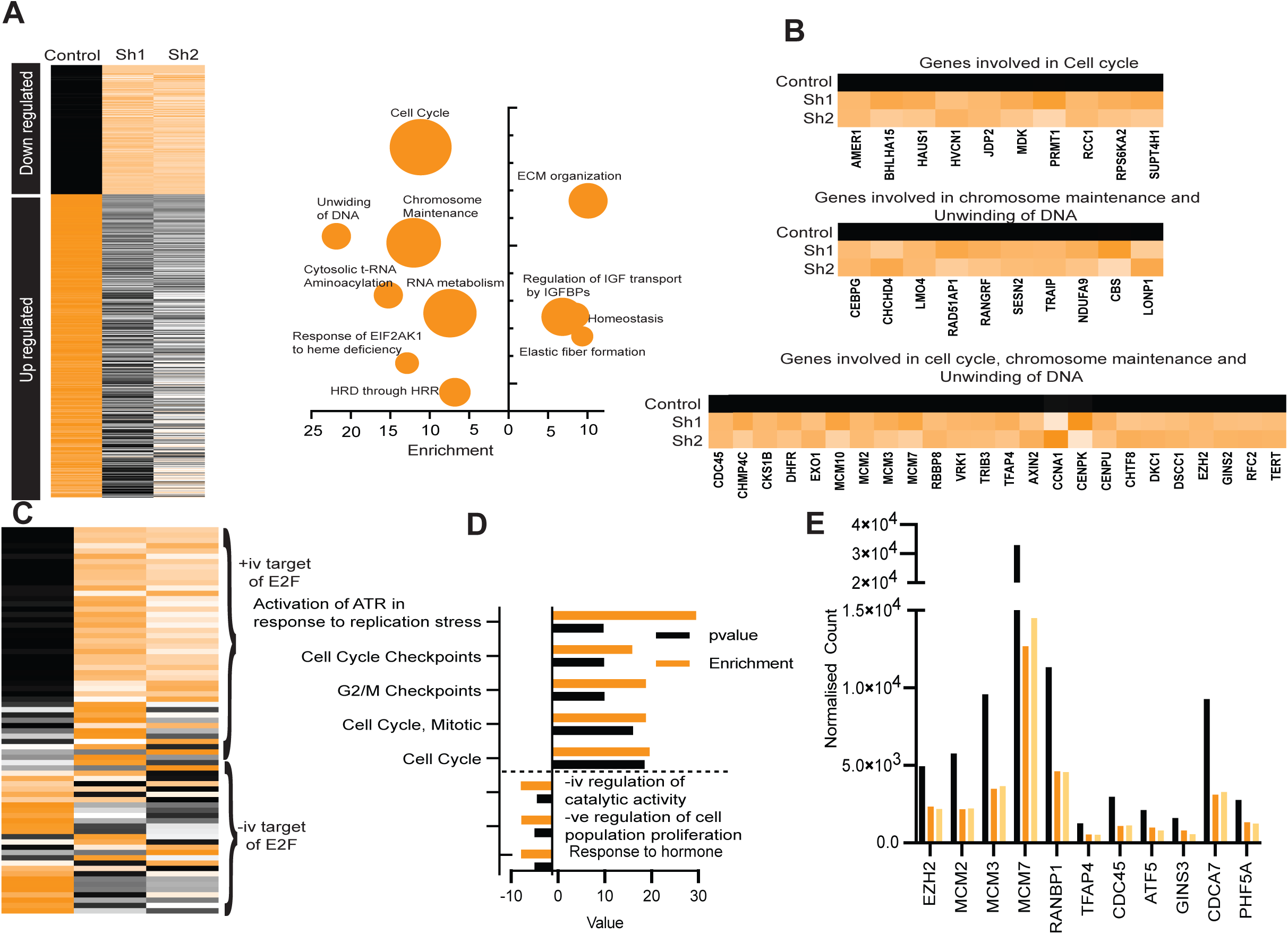
Role of MRPL47 in E2F-Mediated Cell Cycle and Chromosome Dynamics. **(A)** Differential Gene Expression Analysis showing upregulated/downregulated genes in MRL47 knockdown vs control groups. **(B)** Pathway Enrichment Analysis illustrates the major affected pathways and downregulation of Genes Involved in Cell Cycle and DNA Maintenance **(C)** Heatmap depicting the reduced expression of key genes regulating cell cycle progression, chromosome maintenance and DNA unwinding. **(D)** List of positive and negative E2F Target Genes and Checkpoint Regulation **(E)** Functional Enrichment Analysis demonstrating significant impairment in mitotic checkpoints and replication stress response pathways. **(F)** Bar plots reveal a striking reduction in essential DNA replication and cell cycle regulatory genes.

### 3.6 MRPL47 regulates cellular senescence and the ROS-p38MAPK signaling axis

To investigate the mechanism of phosphorylation of Rb increase in MRPL47 knockdown cells, we analyzed p38 phosphorylation, as p38 is a downstream target of ROS. Intriguingly, MRPL47 knockdown decreased ERK1/2 phosphorylation while increasing p38 phosphorylation, suggesting ROS-driven p38 activation. Also, we demonstrated that MRPL47 knockdown suppresses cell growth by inhibiting E2F activity. Since the CDK2-cyclinE complex activates E2F via RB phosphorylation, and its activity is regulated by CDKN1A, we analyzed CDKN1A expression and phospho-Rb levels in MRPL47-depleted cells. Notably, MRPL47 knockdown significantly increased CDKN1A levels, accompanied by a marked reduction in phospho-Rb **(Figure 6A, B and C)** To determine whether elevated reactive oxygen species (ROS) in MRPL47-deficient cells drive phospho-Rb level increase, we treated cells with the ROS scavenger N-acetylcysteine (NAC). Strikingly, NAC restored phospho-Rb levels, confirming ROS-mediated E2F inactivation. **(Figure 6D)** Given that increase in CDKN1A level and decrease in phosphorylation of Rb induces senescence and G1 arrest ^29^, we assessed these phenotypes in MRPL47 knockdown cells. β-galactosidase assays and western blot of β-galactosidase confirmed increased senescence **(Figure 6E, F)**, while qRT-PCR data revealed reduced CDK4, CDK6, and cyclin D levels **(Figure 6G)**. Cell cycle profiling demonstrated a pronounced G1 accumulation and decreased S-phase entry, corroborated by live-cell imaging using the PIP-FUCCI system **(Figure 6 H, I and J)**. Together, these findings establish MRPL47 as a regulator of the CDKN1A-ROS axis, linking its loss to senescence and cell cycle arrest via E2F suppression. In summary, MRPL47 knockdown elevates ROS levels, activating the p38 pathway to upregulate CDKN1A. The CDKN1A induction drives cell growth inhibition via Rb hypophosphorylation and E2F inactivation, ultimately promoting senescence and G1 arrest.

**Figure 6.**
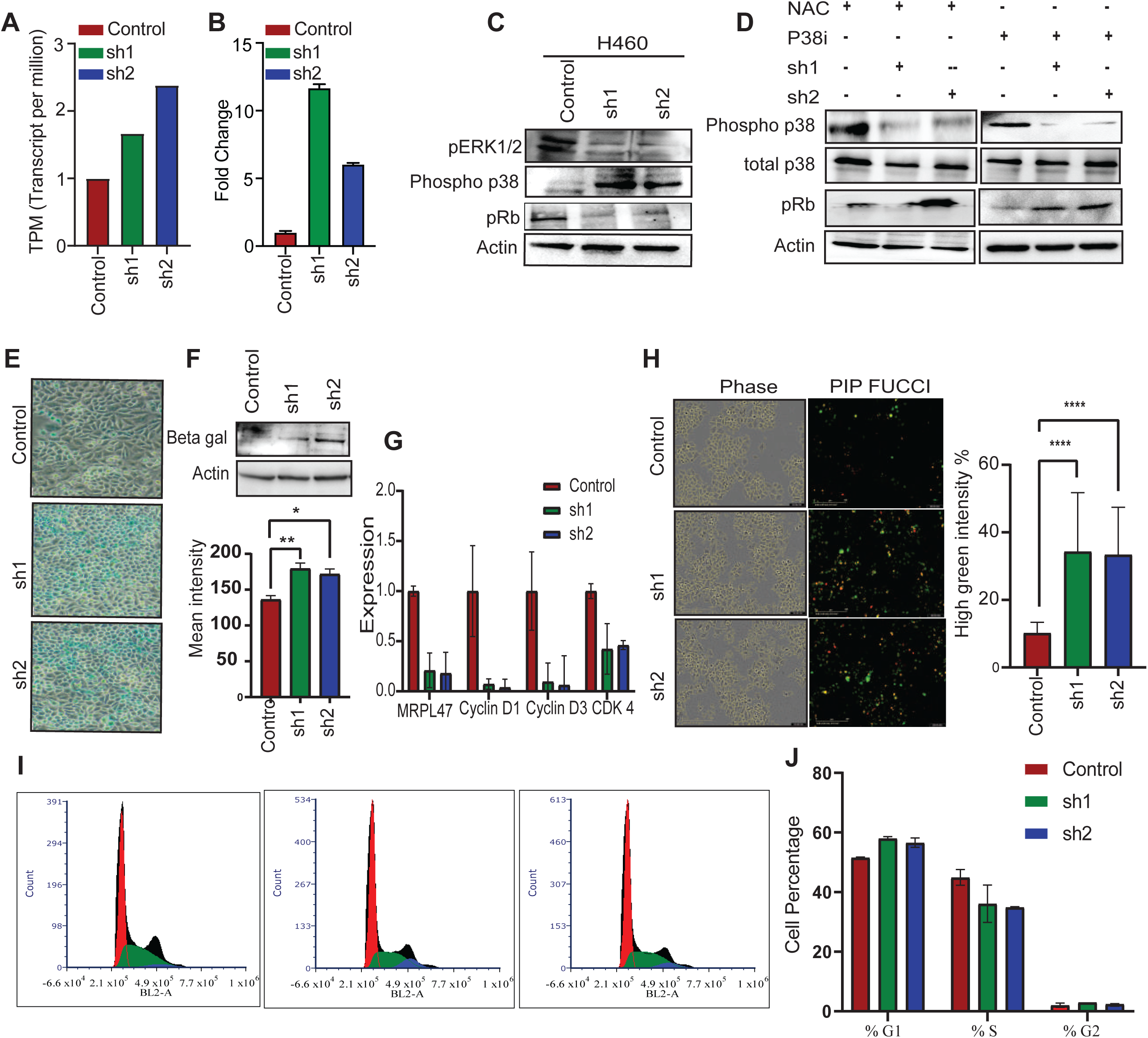
Loss of MRPL47 drives cellular senescence and cell cycle arrest through p38MAPK-CDKN2A Pathway. **(A)** Expression of CDKN1A in control and MRPL47 knock down cells as measure by RNA-seq. **(B)** Expression of CDKN1A in control and MRPL47 knock down cells as measure by real time PCR**. (C)** Western blot showing the phosphorylation levels of key senescence and cell cycle markers including phospho-ERK1/2, phospho-p38 and phospho-Rb in H460 cells. **(D)** Western blot analysis showing the effects of p38 inhibitor (P38i) and antioxidant NAC treatment on p38 phosphorylation, and phospho-Rb levels in MRPL47 knockdown (sh1, sh2) and control H460 cells. **(E)** Beta-galactosidase staining to check senescence in MRPL47 knockdown H460 cells. The blue staining represents senescent cells. **(F)** Western blot analysis of Beta-galactosidase in H460 cells, with quantification of staining intensity (right) showing increased senescence in knockdown cells compared to control. **(G)** qPCRanalysis of Cyclin D1, Cyclin D2 and CDK4 in MRPL47 knockdown. **(H)** Representative images of the PIP-FUCCI reporter assay showing G1 cell cycle arrest (high green intensity) in MRPL47 knockdown. Quantification of high green intensity cells is shown on the right. **(I)** Flow cytometry cell cycle analysis in MRPL47 knockdown. **(J)** The Bar graph shows the percentage of cells in G1, S, and G2 phases.

## 4. Discussion

Mitoribosomes, protein complexes present inside mitochondria plays a piovotal role in translating mitochondrial mRNA into proteins essential vital for oxidative phosphorylation (OXPHOS). Dysregulation of these mitochondrial ribosomal proteins (MRPs) has been associated with cancer progression through alteration in mitochondrial metabolism, impaired oxidative phosphorylation and increased production of ractive reactive oxygen species (ROS), underscoring their therapeutic potential. ^5,30^. We identifed MRPL47 a recurrently amplified and overexpressed gene in NSCLC. Our study delinates the crucial role of MRPL47 in NSCLC tumor progression by regulating mitochondrial function. Loss of MRPL47 compromised mitochondrial translation resulting in decreased expression of selected proteins of Complex I, II and IV subunits of OXPHOS system, thus affecting electron transport chain (ETC) function and consequently increasing mtROS levels. Such malfunction of ETC further lead to altered mitochondrial membrane potential, decreased ATP generation and oxygen consumption rate in the MRPL47 silenced cells. Mechanistically, our findings highlight the link between MRPL47 and mitochondrial dynamics via activation of the p38 MAPK-Rb-E2F pathway resulting in G1 arrest and cellular senescence. These findings underscore MRPL47’s significance in mitochondrial homeostasis and its potential as a therapeutic target in Lung adenocarcinoma.

MRPL47 uniquely influences mitochondrial protein synthesis by selectively regulating translation of ETC subunits of complex I (NDUFB8), II (SDHB) and IV (COX II), while not showing any relation with other proteins of subunits III (UQCRC2) and V (ATP5A). This observation contrasts with conventional perception of mitoribosomes as drivers of global mitochondrial protein synthesis ^31^. This selective regulation of mitochondrial translation by MRPL47 was further confirmed by SUrface SEnsing of Translation (SUnSET) assay, demonstrating that silencing MRPL47 doesn’t impact the synthesis of all mitochondrial-encoded proteins. These findings correspond with recent evidence suggesting that mitochondrial ribosomal proteins (MRPs) may possess distinct functions beyond general translation ^32^. Interestingly, the reduced protein expression of ETC complexes I and III subunits was accompanied with a considerable decrease in their enzymatic activity, highlighting the functional implications of MRPL47 silencing. Since, these complexes are also major contributors of ROS production ^33^ their dysfunction likely contributes as a mechanistic basis for the observed increased in mtROS levels.

Although our findings suggest that MRPL47 may not be directly required for the translation of Complex III subunit protein, its involvement in Complex I translation may indirectly influence Complex III activity.Moreover, the decrease in protein expression of Complex IV subunit (COX II) may be a consequential outcome of compromised mitochondrial translation and malfunction. Given that Complexes I and II are essential for the electron transport chain’s operation, their impairment certainly disturbs the mitochondrial dynamics, restricting co-factor availability and modifying transcription factors such as NRF1 and TFAM ^34^. Furthermore, increased ROS production may also destabilize Complex IV indicating a widespread change in mitochondrial translation and homeostasis, compromising electron transport chain function and mitochondrial efficiency ^35^.

Our findings elucidate the molecular factors influencing the dependency of cancer cells on MRPL47. We comprehensively investigated to demonstrated a strong correlation between dependency on MRPL47 and expression of the tumor suppressor CDKN1A, a well-established regulator of cell cycle progression and senescence. Consequently, an increased CDKN1A protein expression was observed in MRPL47 silenced cells exhibiting growth arrest which was in line with our previous findings. This upregulation was associated with increased phosphorylation of Rb and a decrease in cyclin D and CDK2 levels, leading to G1 cell cycle arrest. These results correspond with earlier studies connecting CDKN1A to the ROS-p38 MAPK pathway in cellular senescence ^36^ and mitochondrial dysfunction ^37^.

A critical aspect of our study is to understand how MRPL47 play role in regulating mtROS signalling. The increased level of mtROS levels in MRPL47 silenced cells were found to be associated with altered mitochondrial membrane potential (ΔΨM), lower oxygen consumption rates and reduced ATP synthesis collectively signifying mitochondrial dysfunction. Furthermore, to understand whether ROS is mediating the cell growth inhibition in the silenced cells, NAC treatment was provided to scavenge ROS. The cell growth was rescued, confirming that ROS mediates the phenotypic consequences of MRPL47 knockdown. The downstream effects of elevated mtROS were notably apparent in the activation of the p38 MAPK pathway, a key modulator of cellular stress responses and carcinogenesis ^38^. The phosphorylation of p38 and its downstream target, Rb, indicates the role of ROS-p38-MAPK signalling in the regulation of cell cycle arrest. Additionally, RNA sequencing analysis further confirmed that MRPL47 inhibits cell growth by downregulating E2F, a key regulator of cell cycle, showing consistency with our prior findings. These findings highlight the association between MRPL47, ROS signalling, and cell cycle regulation.

The therapeutic significance of MRPL47 was highlighted by its amplification and overexpression in several cancer types, especially non-small cell lung cancer (NSCLC). Analysis of TCGA data demonstrated that MRPL47 amplification is substantially correlated with its expression, aligning with prior research indicating that gene amplification serves as a catalyst for oncogene overexpression ^39^. In LUAD patients, elevated MRPL47 expression correlated with poor prognosis, reinforcing its potential as a prognostic biomarker.

Our findings illustrate that Mitoribosomes have a role beyond their canonical functions in mitochondrial translation, uncovering their selective translational activity and influence on ROS signalling. Subsequent research delineates the domain of MRPL47’s substrate’s specificity through methods like cryo-electron microscopy which will provide an understanding of its unusual form of its distinct translational control. In vivo investigations are crucial to confirm the therapeutic efficacy of targeting MRPL47 in cancer models. Given the relationship between mitochondrial dysfunction and the tumor environment, MRPL47 mediated ROS generation’s effect on immune cell recruitment and activation is crucial to investigate. Such research may reveal innovative combinatorial approaches utilising MRPL47 inhibitors alongside immunotherapies.

## Acknowledgements

Sudhanshu Shukla would like to acknowledge Indian Council for medical research, Govt. of India (Grant # 2021-9513/CMB/ADHOC-BMS and EMDR/SG/13/2023-0244), Department of Biotechnology, Govt of India (Grant id – BT/PR51308/MED/30/2511/2023) and Anusandhan National Research foundation, Govt. Of India (Grant ID – CRG/2023/000837 for the funding. LLM based AI databases were used for the paraphrasing and Grammer correction.

## Author contributions

**NB:** Conceptualization, Methodology, Formal analysis, Investigation, review and editing.

**DA**: Investigation

**AC**: Investigation

**SY:** Investigation

VB: Investigation

**BKC**: Review and editing, Supervision

**SS**: Conceptualization, Methodology, Formal analysis, Writing – Review & Editing, Funding Acquisition.

## Figure Legend

**Supplementary Figure 1:**
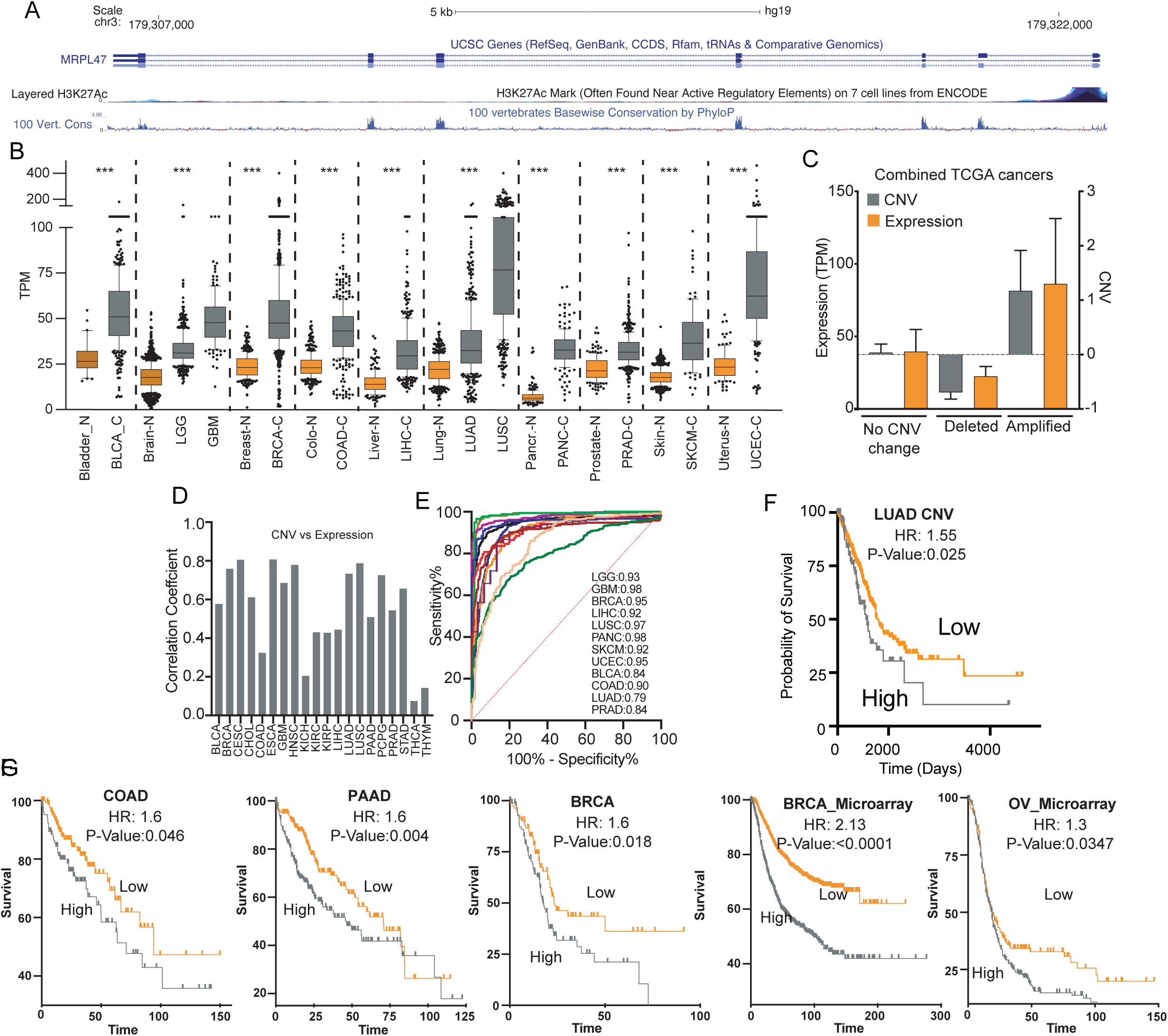
**A)** Structure of MRPL47 gene. Chip-seq data showing enrichemnt of H3K27Ac. **B**) Expression of MRPL47 RNA across multiple common cancer and correponding normal samples. **C**) Bar chart showing the CNV and expression association of MRPL47 gene in all TCGA cancer types. **D**) Correlation between expression and CNV was calculated and plotted to show the high association of MRPL47 expression and CNV. **E**) ROC plots showing the cancer specific expression of MRPL47. F) Prognostic association of MRPL47 CNV in LUAD **G**) Prognostic role of MRPL47 expression in given cancer types.

